# Pharmacological inhibition of all known major inward cationic currents does not block the induction of spreading depolarizations

**DOI:** 10.1101/2025.07.16.665203

**Authors:** Preston C. Withers, Allen Jones, Kojo Bawuah Afran-Okese, Bailey Calder, Hunter J. Morrill, T. Luke Shafer, Dallin S. Nevers, Jacob Norby, Rebeca Acosta, Benjamin T. Bikman, Arminda Suli, R. Ryley Parrish

## Abstract

Spreading depolarization (SD) is a wave of profound cellular depolarization that propagates across central nervous system tissue and causes a near-complete collapse of ionic gradients. Implicated in neuropathologies including seizures, migraine with aura, traumatic brain injury, and stroke, SD is experimentally induced in animals by electrical stimulation, mechanical injury, hypoxia, elevated extracellular potassium, and various other techniques. Despite extensive research, the mechanisms underlying SD initiation remain unclear. Prior research in rodents found that simultaneously blocking sodium, calcium, and glutamatergic (AMPA and NMDA) channels prevents SD induction whereas inhibiting any two of these three currents is insufficient. This suggests that SD induction could be a product of overstimulation of any single known inward cationic current. However, some researchers propose that SD induction occurs via an unknown “SD channel.” To further explore the role of known inward cationic currents in SD induction, we applied high potassium to two biological models, namely zebrafish and mice. First, we developed a novel *ex vivo* zebrafish model to assess SD induction in the optic tectum. Using KCl microinjection and DC local field potential recordings, we found that inhibition of sodium, calcium, and glutamatergic channels significantly decreased SD amplitude but never blocked SD induction in the zebrafish optic tectum. Similar pharmacological experiments in hippocampal mouse slices (CA1 subregion) also confirmed that SDs persist despite the same pharmacological cocktail. These findings suggest that additional mechanisms beyond sodium, calcium, and glutamatergic signaling contribute to SD induction, supporting the hypothesis that a currently unknown channel is critical in SD physiology.

## Introduction

Spreading depolarization (SD) is a complex and poorly understood event characterized by a wave of extreme neuronal depolarization which travels through central nervous system tissue and disrupts normal ionic gradients. SD was first identified by Aristides Leão in 1944 as a slow, propagating wave that suppressed epileptiform activity, hence the near-synonymous term “spreading depression” (Leao, 1947). SDs are implicated in numerous neuropathologies, including seizures (Aiba et al., 2023; Norby et al., 2025; Tamim et al., 2021), migraines (Chever et al., 2021), traumatic brain injuries (Mosley et al., 2022), subarachnoid hemorrhage (Nasretdinov et al., 2023; Zheng et al., 2020), and ischemic strokes (Binder et al., 2022; Torteli et al., 2023). Although recognized as a key contributor in human disease, how SDs are induced remains an enigma.

SD can be induced experimentally by a variety of methods that disrupt ionic homeostasis (Aiba et al., 2023; Binder et al., 2022; Chever et al., 2021; Herreras & Somjen, 1993; Juzekaeva et al., 2020; Mosley et al., 2022; Nasretdinov et al., 2023; Tamim et al., 2021; Torteli et al., 2023; Zheng et al., 2020). Previous findings suggest that inhibiting sodium, calcium, and glutamatergic currents can modulate or altogether prevent experimentally induced SDs, highlighting the role of these currents in SD induction (Alday et al., 2014; Klass et al., 2018; McLachlan, 1992; Rashidy-Pour et al., 1995; Vitale et al., 2023; Wang et al., 2012). For instance, prior research in rats asserts that blocking sodium, calcium, and glutamatergic (AMPA and NMDA) channels completely prevents SD induction (Muller & Somjen, 1998), while inhibiting just one or two of these currents is insufficient to block SD. This suggests that any of these currents is sufficient to initiate an SD (Muller & Somjen, 1998).

SDs, therefore, may be a non-specific event induced by various pathways, provided there is sufficient cellular stress and residual cationic inward current, which results in strong neuronal depolarization. Contrary to the mentioned hypothesis, some have proposed the existence of an SD-specific transmembrane protein (Aitken et al., 1998; Basarsky et al., 1999; Chen et al., 2017; Ricks et al., 2025; Wang et al., 2024). Candidates include pannexins (Chen et al., 2017), volume-regulated anion channels (VRACs) (Basarsky et al., 1999; Ricks et al., 2025), gap junctions (Aitken et al., 1998; Kunkler & Kraig, 1998), and the Na/K ATP-ase (Kim et al., 2025; Wang et al., 2024), all of which could facilitate network depolarization and alter cellular swelling, complicating the potential mechanisms behind SD induction.

SDs have been found to be evolutionarily conserved across mammals, amphibians, fish, and insects (Dell’Orco et al., 2023; Spong et al., 2016; Terai et al., 2021), allowing for the expansion of SD research beyond the limitations of traditional mammalian systems. Notably, non-mammalian models offer greater affordability, rapid development cycles for more efficient genetic manipulation, and suitability for high throughput screening, potentially accelerating discovery in SD research (Spong et al., 2016; Terai et al., 2021). Although insect models show promise in SD research, key physiological differences from mammals, including invertebrate status, resistance to oxygen glucose deprivation and toxin treatments, and distinct ion regulatory mechanisms, complicate comparisons to mammals (Robertson et al., 2020; Robertson & Wang, 2025; Wang et al., 2024). To overcome these challenges, a recent *in vivo* study used zebrafish (*Danio rerio*) as an alternative model, with initial characterization demonstrating similar SD propagation rates, recovery times, and amplitudes to those observed in rodent models (Terai et al., 2021). Zebrafish are well-suited vertebrates for studying ion channel dynamics due to their genetic tractability, optical transparency, and cost-effective high-throughput screening potential (Alday et al., 2014; Cozzolino et al., 2020; Cully, 2019; Dixon et al., 2023; Kimmel et al., 1995). Despite these advantages, the zebrafish model remains underutilized for uncovering the cellular basis of SD.

To further clarify the mechanisms of SD initiation while taking advantage of the ease of channel antagonist/agonist application and improved control and replicability of *ex vivo* models, we developed a whole-brain *ex vivo* zebrafish protocol to characterize the contribution of known inward cationic currents to SD induction. Contrary to previous rodent findings (Muller & Somjen, 1998), inhibiting sodium, calcium, and glutamatergic channels did not block SDs in the zebrafish optic tectum following KCl microinjection. Confirming these results using an analogous protocol in mouse hippocampal slices, we again found that inhibiting sodium, calcium, and glutamatergic channels with TTX, MK-801 (or APV), NBQX, and NiCl_2_ did not block SDs. However, this cocktail of cationic current inhibitors was able to reduce SD amplitude in our novel zebrafish model. Collectively, these findings suggest that, while known inward currents may modulate total SD current, they are not solely responsible for SD induction, strengthening the hypothesis that additional players are critical to SD physiology.

## Methods and Materials

### Zebrafish Brain Preparations

All animal procedures were performed in accordance with regulations of the Institutional Animal Care Committee and the Brigham Young University Animal Care Committee. Male and female adult zebrafish (AB strain, 3-6 months old, n=18) were used. A subset (n=10) had been injected as larvae with a halorhodopsin plasmid under either the UAS or huc promoters. With all fish used in these experiments, SDs were not blocked when exposed to the inhibitory compounds (n=18), and no significant difference in the SD amplitude percent change was found between the two fish groups (AB strain or the halorhodopsin injected fish) when the full cocktail was added (unpaired t test, n=12, p=0.5905). Therefore, the two groups were combined and reported together. Fish were kept on a 14 h:10 h light:dark cycle, at 28.5°C, and fed brine shrimp. Fish were anesthetized using 0.04% MS222 (tricaine mesylate, pH 7.2) until immobile and unresponsive. Zebrafish were placed on filter paper (P4, Fisherbrand) and decapitated posterior to the gills to ensure a complete brain sample. Zebrafish heads were placed in a pre-cut sponge groove in a petri dish secured with 2 insect pins in carbogenated (95% O2 and 5% CO2) artificial cerebrospinal fluid (aCSF) containing the following (in mM): 1 MgCl2; 2 CaCl2; 126 NaCl; 26 NaHCO3; 3.5 KCl; 1.26 NaH2PO4; 10 glucose. Under a SZ61 microscope (Olympus), a small midline incision was made at the base of the skull, and the skull hemispheres were carefully removed using forceps. The optic nerves were severed by lifting the brain with forceps and cutting them with dissection scissors (Figure 1A). Following dissection, zebrafish brains were placed in a custom holding chamber (Withers et al., 2024) with aCSF and allowed to rest for 30 minutes prior to experimentation.

**Figure 1.**
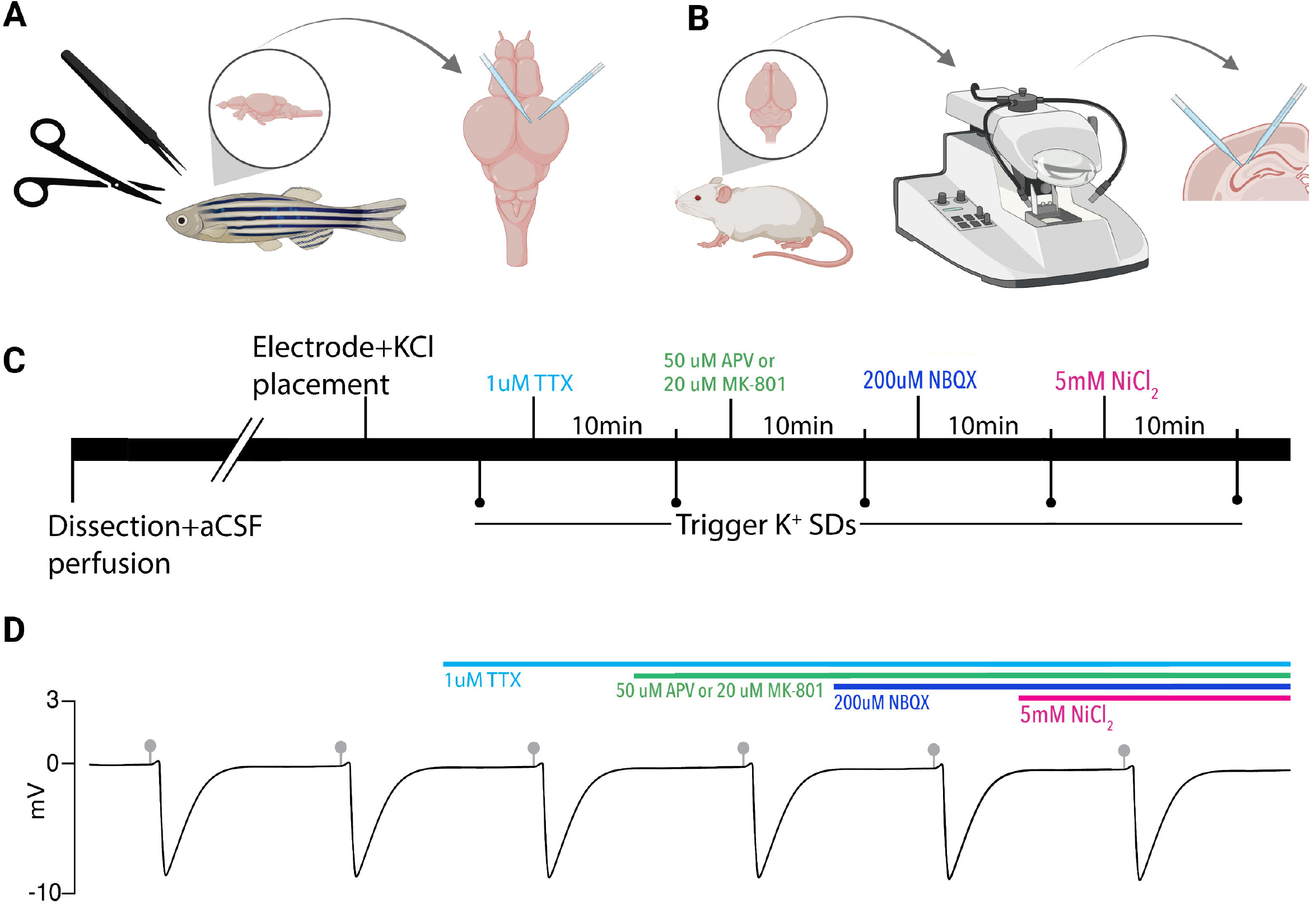
Experimental paradigm. (A) Zebrafish brains were removed and electrodes were placed in the optic tectum for SD recordings and KCl microinjection. (B) Mouse brains were removed, sectioned on a vibratome, and electrodes were placed in the CA1 region of the hippocampus for recording of SDs and KCl microinjection. (C) Timeline of pharmacological application used in both models. SDs were evoked via picospritzer 10 minutes after each compound application. (D) A model DC LFP trace showing persistent SDs induced by KCl microinjections (gray markers). Colored bars indicate drug application timing. Created in BioRender. https://BioRender.com/8sngb4d.

### Mouse Brain Slice Preparations

All animal procedures were performed in accordance with the Institutional Animal Care Committee and Brigham Young University Animal Care Committee. Experiments were performed on male and female C57BL/6 mice (4–12 weeks old). Mice were housed in individually ventilated cages in a 12:12 h light:dark cycle. Mice were anesthetized with isoflurane, decapitated, and had their brains harvested. Brains were placed in cold cutting solution perfused with carbogen and containing the following (in mM): 3 MgCl_2_; 126 NaCl; 26 NaHCO_3_; 3.5 KCl; 1.26 NaH_2_PO_4_; 10 glucose. 350 μm horizontal brain slices were made on a Leica VT1200 vibratome (Leica Microsystem) and further dissected into hippocampal sections with a scalpel (Figure 1B). Slices were stored for 1-4 hours at room temperature in a custom-made holding chamber before experimentation. Holding chamber solutions were bubbled with carbogen (95% O2 and 5% CO2) in aCSF containing (in mM): 2 CaCl_2_; 1 MgCl_2_; 126 NaCl; 26 NaHCO_3_; 3.5 KCl; 1.26 NaH_2_PO_4_; 10 glucose.

### Recording Solutions

aCSF contained either (in mM): 2 CaCl_2_; 1 MgCl_2_; 126 NaCl; 26 NaHCO_3_; 3.5 KCl; 1.26 NaH_2_PO_4_; 10 glucose, or 1.2 CaCl_2_; 1.2 MgCl_2_; 130 NaCl; 24 NaHCO_3_; 3.5 KCl; 10 glucose in these studies. NaH_2_PO_4_ was excluded in the second aCSF to ensure NiCl_2_ did not precipitate out of solution, which has been suggested to be a problem in prior studies (Muller & Somjen, 1998). To control for this potential confounding factor, four zebrafish experiments and all mouse slice experiments used aCSF without NaH_2_PO_4_. Both aCSF recipes demonstrated similar results in the zebrafish, and no significant difference in the SD amplitude percent change was observed between the two recipes with the addition of the full cocktail (unpaired t test, n=12, p=0.7615), so the data were combined. All aCSF solutions were bubbled with carbogen (95% O2, 5% CO2).

Compounds were dissolved in water and used at the following concentrations: TTX (tetrodotoxin; Hello Bio) was used at 1 µM, MK-801 (Dizocilpine; Hello Bio) at 20 µM, NBQX (2,3-dioxo-6-nitro-7-sulfamoyl-benzo[f]quinoxaline; Hello Bio) at 20 µM, APV (also known as AP5, (2R)-amino-5-phosphonovaleric acid, or (2R)-amino-5-phosphonopentanoate; Hello Bio) at 50 µM, and NiCl_2_ (Thermo Fisher Scientific) at 5mM.

### Zebrafish Electrophysiology Recordings

The brains were submerged in a custom 3D-printed recording chamber, secured with a platinum horse-hair harp, and continuously perfused with ∼36°C aCSF at 4mL/min using a peristaltic perfusion system (PPS2 Multi Channel Systems) and an inline heater (Solution Heater SH-27B, Warner Instruments). Recordings were obtained using aCSF-filled ∼5 MΩ borosilicate glass micropipettes (GC120TF-10; Harvard Apparatus) and the same micropipettes were used for KCl microinjection. Both micropipettes were placed in the optic tectum using micromanipulators (Scientifica and Siskiyou respectively). Electrode tips were placed adjacently and 80-100µm deep in the optic tectum (Figure 1A). Spreading depolarizations (SDs) were induced by microinjecting 100mM KCl with a picospritzer III system (Parker Hannifin Corporation) set to 30 psi. DC local field potential (LFP) recordings were acquired using Clampex with a CV-7B Current Clamp and Voltage Clamp Headstage (Axon Instruments), amplified 10 times with a Multiclamp 700B amplifier (Axon Instruments), and digitized with a Digidata 1440A analog-to-digital converter (Axon Instruments). Before adding inhibitors, SD inducibility was confirmed by at least two SDs separated by 10 minutes of rest. Inhibitors were added sequentially to the aCSF bath to obtain the desired concentrations with a 10-minute wait time between drug addition and KCl microinjection. Inhibitors were applied in varying orders of application to analyze their individual and collective effects on KCl-induced SDs (Figure 1C–D). MK-801 has previously been reported to block SD induction, so we performed experiments using either APV or MK-801 to block NMDA receptor-mediated currents (Anderson & Andrew, 2002; Vitale et al., 2023) (Figure 1C–D). Compounds were applied sequentially in the following orders: TTX, NiCl_2_, APV, and NBQX (n=1); TTX, NiCl_2_, and then APV and NBQX simultaneously (n=5); TTX, MK-801, NBQX, and NiCl_2_ (n=3, Figure 1C–D); TTX, NiCl_2_, MK-801, and NBQX (n=2); and TTX, NiCl_2_, MK-801, and NBQX simultaneously (n=1). The complete combination of cationic current inhibitors, including TTX, NBQX, NiCl_2_, and either APV or MK-801, was collectively designated as the “full cocktail.”

### Mouse Electrophysiology Recordings

The slices were transferred to a custom-made submersion chamber (Withers et al., 2024), secured with a platinum horse-hair harp, and perfused with ∼36°C aCSF at 4.3 mL/min using a peristaltic perfusion system (PPS2 Multi Channel Systems) and an inline heater (SH-27B Harvard Apparatus and TC-344B, Warner Instrument Corporation). Recordings were obtained using aCSF-filled ∼1 MΩ borosilicate glass micropipettes (GC120TF-10; Harvard apparatus) and positioned ∼150 µm deep in the CA1 pyramidal layer for LFP recordings, recorded using Wavesurfer 1.0.6 via MATLAB 2024b (Figure 1B). ∼5 MΩ glass micropipettes were backfilled with 1 M KCl and secured in a headstage, connected to a picospritzer II system (Parker Hannifin Corporation), then positioned in the slice near the recording electrode at the same depth. SD events were induced using KCl microinjection at 30 psi regulated by the picospritzer. The DC signal was recorded at 10 kHz, amplified 10X using a custom-built amplifier, and digitized with an NI USB-6341 X Series Multifunction DAQ board (National Instruments). Compounds were bath applied sequentially in the following order: TTX, APV, NBQX, and NiCl_2_ (n=2, Figure 1C–D). MK-801 has been hypothesized to block SD induction in prior studies, so we performed an additional experiment with the same cocktail order, only replacing APV with MK-801 (n=1, Figure 1C–D).

### Data Analysis

Electrophysiology recordings were analyzed using custom MATLAB programs (available upon request). The negative DC voltage shift amplitude and duration following KCl microinjection were documented for each SD in both animal models with and without the full pharmacological cocktail. SDs were considered blocked if consecutive KCl microinjection-induced SD events ceased following drug application and returned after aCSF washout. For each zebrafish brain and mouse brain slice, SD amplitudes were averaged within each pharmacological condition. Mean SD amplitude under control conditions (without inhibitors) was compared to the mean amplitude under each drug combination. The percent change between paired means was calculated with standard error. Group SD amplitude mean and standard error were also computed across all zebrafish brains and mouse slices for each inhibitory condition.

Standard errors were not reported for conditions containing only one or two samples. A Wilcoxon matched-pairs signed-rank test was performed on the mouse and zebrafish data to assess differences in average SD amplitudes and durations before and after pharmacological blockade of all three major inward currents (full cocktail containing either MK-801 or APV, Figure 2 C–D, H–I). SD duration was defined as the time interval between the onset of the negative voltage shift and the point at which the signal returned to 50% of baseline (Terai, 2021). Matched pairs consisted of SDs recorded from the same zebrafish brain or mouse brain slice with or without pharmacological solutions. Wilcoxon tests and normality tests were performed using BioRender, and paired t-tests were performed using custom MATLAB code. Figures were created using BioRender (accessed July 2025) under a publication license (Premium Academic plan), with the specific figure URLs provided where applicable.

**Figure 2.**
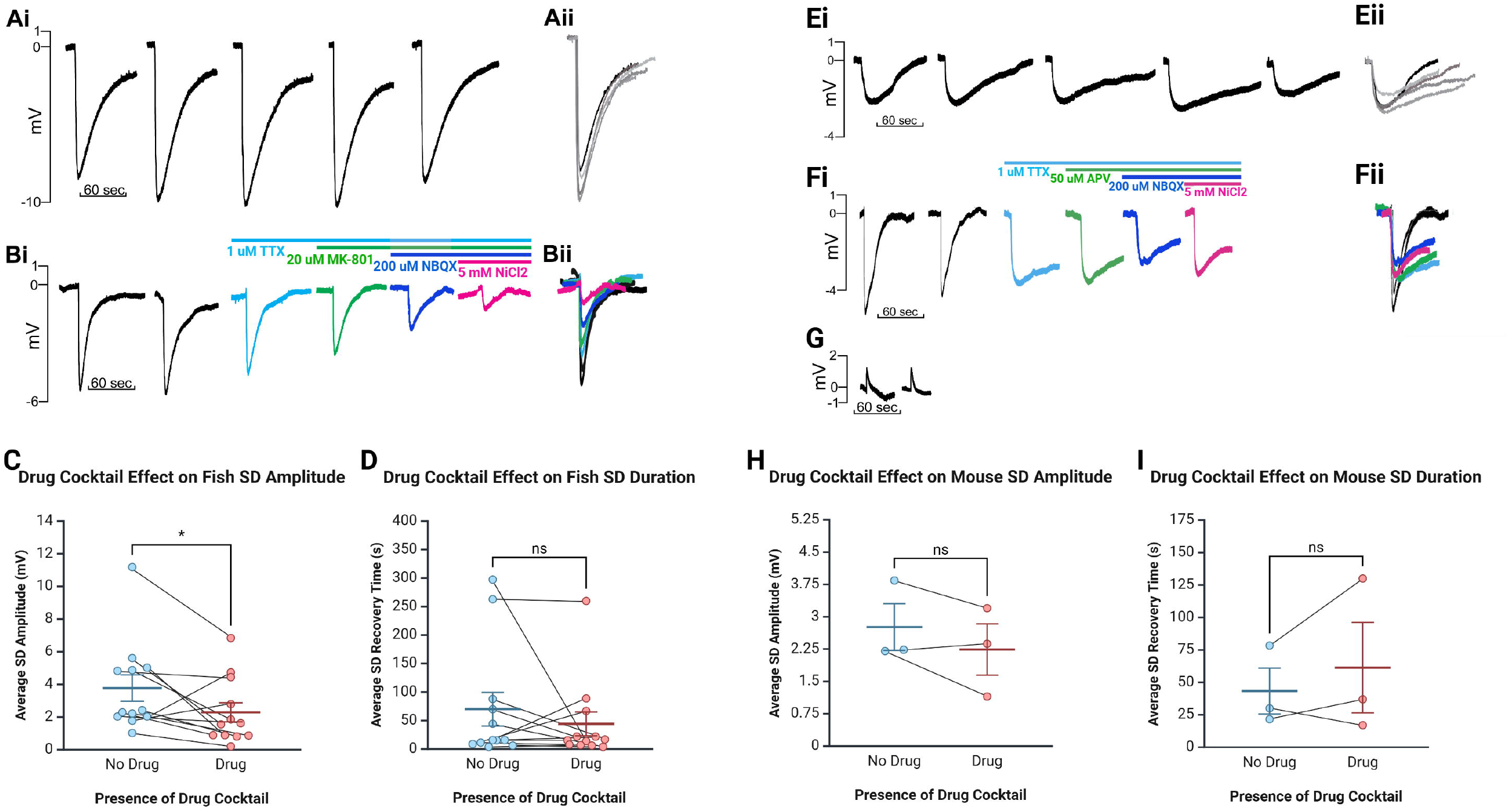
Major inward cationic channel inhibitors significantly decrease SD amplitude in the zebrafish *ex vivo* optic tectum and show a trend toward reduction in mouse hippocampal slices but do not significantly impact SD duration in either model. (Ai-Bii) Representative DC LFP traces (Ai, Bi) and overlays (Aii, Bii) of KCl-induced SDs in *ex vivo* zebrafish optic tectum without compounds (Ai-Aii) and following stepwise drug application (Bi-Bii). (C-D) SD amplitude in the zebrafish model was significantly decreased following the full pharmacological cocktail (Wilcoxon matched-pairs signed-rank test, W=54, n=12, p=0.0342), while the duration did not significantly change (Wilcoxon matched-pairs signed-rank test, W=19, n=12, p=0.533). (Ei-Fii) Representative traces (Ei, Fi) and overlays (Eii, Fii) of SDs in mouse CA1 without compounds (Ei) and with progressive compound application (Fi). (G) Representative trace of a failed SD induction after KCl injection. (H-I) In mouse hippocampal slices, changes in SD amplitude (H, Wilcoxon matched-pairs signed-rank test, W=4, n=3, p=0.5) and duration (I, Wilcoxon matched-pairs signed-rank test, W=-4, n=3, p=0.5) with the addition of the cocktail were insignificant. Box-and-whisker plots show mean ± SEM. Inhibitory cocktail included TTX, MK-801 or APV, NBQX, and NiCl□. Created in BioRender. https://BioRender.com/ttlns2v.

## Results

### Ex vivo zebrafish optic tectum as a model of spreading depolarizations

Spreading depolarizations (SDs) are evolutionarily conserved and inducible from insects to humans (Norby et al., 2025; Robertson et al., 2020; Robertson & Wang, 2025; Wang et al., 2024). Although an *in vivo* zebrafish SD model exists (Terai et al., 2021), an *ex vivo* model, which would increase the ease of channel manipulation and improve experimental control and reproducibility, has yet to be developed (Hamilton & Santhakumar, 2020; Vaughan et al., 2024). Therefore, in this study, we aimed to determine if an *ex vivo* zebrafish brain could generate SDs and serve as a model for studying SD induction. Using dissected zebrafish brains and recording from the optic tectum (Figure 1A), we found that these preparations reliably produced characteristic, repetitive SDs when microinjected with KCl (Table 1), similar to rodent *ex vivo* preparations (Anderson & Andrew, 2002). A distinct advantage of this model compared to most *ex vivo* rodent models is that these brains are small enough to be used intact, without requiring brain sectioning for experimentation, resulting in a whole-brain *ex vivo* preparation.

**Table 1.**
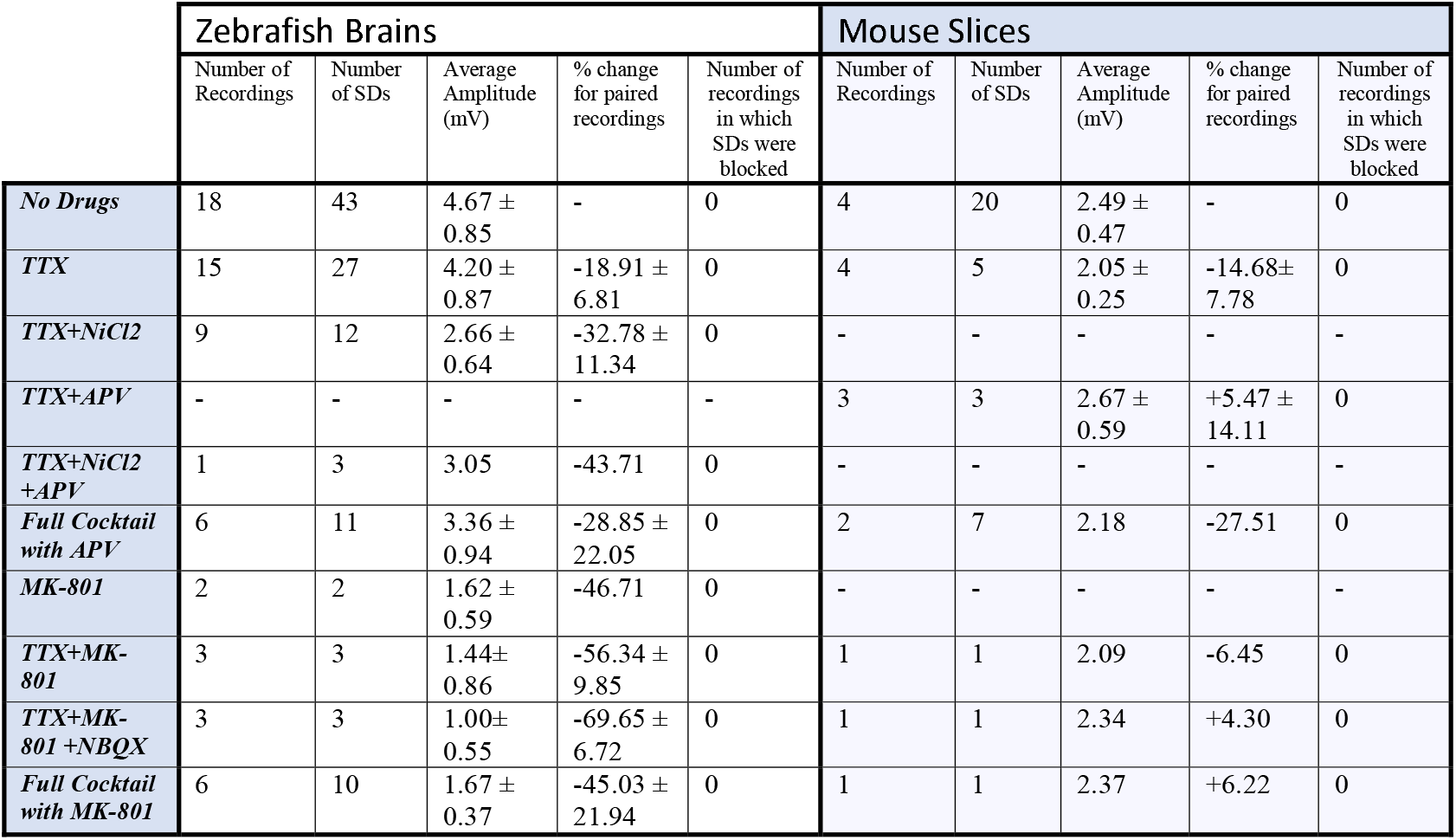
Major inward cationic channel inhibitors do not block SDs in *ex vivo* zebrafish or mouse hippocampal slice preparations but decrease SD amplitude in zebrafish. Each paired recording used to calculate the change in SD amplitude includes a sample in aCSF and the corresponding compounds listed. The symbol ± denotes the standard error of the mean (SEM). SD amplitudes without compounds were only included if obtained during experiments in which at least one compound was used. The “full cocktail with APV” consisted of 1µM TTX, 20µM NBQX, 5mM NiCl_2_, and 50µM APV. The “full cocktail with MK-801” consisted of 1µM TTX, 20µM NBQX, 5mM NiCl_2_, and 20µM MK-801. The number of recordings varied across regimens and not all recordings were continued to the full cocktail.

### Inhibition of major inward ion currents does not prevent SD induction

After confirming that the *ex vivo* zebrafish brain is a reliable model for SD induction, we examined whether SDs generated in this model via KCl microinjection could be blocked by compounds previously demonstrated to inhibit SDs (Anderson & Andrew, 2002; Somjen, 2001; Somjen, 2004; Vitale et al., 2023). Only zebrafish brains that could generate at least 2 SDs were used for pharmacology studies. Once this was confirmed, we bath applied TTX to block voltage-gated sodium channels (Figure 1C–D). As expected, SDs persisted, but their amplitude was reduced based on paired recordings (paired t-test, n=15, p=0.0263, 5.10 ± 0.98 mV before TTX application, 4.20 ± 0.87 mV after TTX application, Table 1). The addition of NiCl_2_ (to block calcium currents) to the TTX solution did not block SDs but reduced SD amplitude further in paired recordings in a trend nearing significance (paired t-test, n=9, p=0.0690, 3.77 ± 0.88 mV before NiCl_2_ application, 2.66 ± 0.64 mV after NiCl_2_ application, Table 1). Additionally, adding MK-801 (to block NMDA currents) to the TTX did not block SDs but reduced the amplitude in each case (paired t-test, n=3, p=0.223, 1.86 ± 1.12 mV before MK-801 application, 1.44 ± 0.86 mV after MK-801 application, Table 1). In a final subset of experiments, additional compounds were added until the zebrafish brains were exposed to TTX, NiCl_2_, NBQX (kainate and AMPA receptor inhibitor), and either APV (NMDA receptor inhibitor) (n=6) or MK-801 (n=6). We refer to this combination of channel inhibitors (Na□ channels, Ca^2^□ channels, and kainate, AMPA, and NMDA receptors) as the “full cocktail.” Despite inhibiting all known inward cationic currents with this pharmacological mixture, SDs persisted with both the full cocktail containing APV (n=6, 5.23 ± 1.30 mV before adding any compounds, 3.36 ± 0.94 mV with full cocktail, Table 1) and the full cocktail containing MK-801 (n=6, 2.45 ± 0.55 mV before adding any compounds, 1.67 ± 0.37 mV with full cocktail, Table 1). However, the full cocktail with either APV or MK-801 did significantly reduce SD amplitude (Figure 2Ai–C, Wilcoxon matched-pairs signed-rank test, W=54, n=12, p=0.0342). We did not find that the SD duration, defined as the time between the initial negative voltage shift and the return to 50% baseline, significantly changed in the presence of the full cocktail (Figure 2D, Wilcoxon matched-pairs signed-rank test, W=19, n=12, p=0.533), despite a reduction in SD amplitude. These data suggest that, although many cationic currents contribute to SD size, voltage-gated Na^+^ channels, voltage-gated Ca^2+^ channels, and glutamatergic receptor-mediated channels (kainate, AMPA, and NMDA receptors) are not responsible for SD induciton in the zebrafish optic tectum.

Because the *ex vivo* zebrafish optic tectum is a novel preparation for studying SDs, we confirmed these results in mouse hippocampal brain slices (Figure 1B–D), a well-characterized and highly susceptible mammalian preparation for SDs (Aiba, 2024; Andrew et al., 2022; Frank et al., 2024; Somjen, 2001; Sword et al., 2024). As in the zebrafish, the full cocktail again did not block KCl-induced SDs. However, unlike in zebrafish preparations, SD amplitude changes were not significant (Wilcoxon matched-pairs signed-rank test, W=4, n=3, p=0.5, Figure 2Ei–H, see also Table 1). The capacity for SD generation was again preserved regardless of the NMDA receptor inhibitor (MK-801 or APV, Table 1) used, and SD duration remained unchanged (Wilcoxon matched-pairs signed-rank test, W=-4, n=3, p=0.5). The rodent data confirm our zebrafish optic tectum findings, indicating that the simultaneous inhibition of all major inward cationic currents is insufficient to block KCl-induced SDs.

## Discussion

Spreading depolarizations (SDs) are a known phenomenon in a number of neurological disorders such as stroke (Binder et al., 2022; Torteli et al., 2023), migraine with aura (Chever et al., 2021), traumatic brain injury (Mosley et al., 2022; Nasretdinov et al., 2023; Zheng et al., 2020), and epilepsy (Aiba et al., 2023; Norby et al., 2025; Tamim et al., 2021). Despite these known contributions to various neurological disorders, the biophysical mechanisms underlying SD induction remain unclear. Here, we developed an *ex vivo* zebrafish optic tectum preparation to investigate the cellular and ionic mechanisms underlying SD. This model reliably produced characteristic SDs in response to high-potassium microinjection, with electrophysiological features (e.g., amplitude, recovery dynamics) comparable to rodent models (Kim et al., 2025; Terai et al., 2021). To our knowledge, this is the first demonstration of SDs in an isolated zebrafish brain, building upon earlier *in vivo* work (Terai et al., 2021) and offering an accessible, easy-to-use, inexpensive, and genetically tractable platform for SD research (Alday et al., 2014; Cozzolino et al., 2020; Cully, 2019; Dixon et al., 2023; Kimmel et al., 1995).

Using this zebrafish preparation, our primary objective was to determine whether SDs could be induced after blocking all known major inward cationic currents. Notably, even after pharmacologically blocking voltage-gated sodium (TTX), calcium (NiCl□), and glutamate (NBQX, APV or MK-801) channels, SD induction remained. We confirmed this finding in a mouse hippocampal slice model, where the same cocktail also failed to prevent SDs. Although SD amplitude was reduced in zebrafish, SD persistence suggests that these widely accepted conductances are not individually or collectively required for SD initiation in the high-potassium model. These findings challenge previous reports that SDs cannot occur when all these conductances are blocked (Muller & Somjen, 1998; Somjen, 2004). While these prior reports did not claim SDs are triggered by these known channels (voltage-gated Na^+^ channels, voltage-gated Ca^2+^ channels, and glutamatergic receptors), they implied that at least one must be recruited for SD induction. The discrepancy may stem from differences in SD induction models: Muller et al., 1998 used hypoxia-induced SDs, whereas we used high-potassium induction. Nevertheless, our findings suggest that SDs can occur despite this full pharmacological blockade, pointing to biophysical changes that could result due to ionic imbalance following KCl injection, creating an osmotic crisis for neurons, triggering SDs through a still unknown process (Parrish, Jackson-Taylor, et al., 2023; Parrish, MacKenzie-Gray Scott, et al., 2023; Ricks et al., 2025). However, reduced SD amplitudes from our study and others (Muller & Somjen, 1998; Suryavanshi et al., 2022; Wang et al., 2012) indicate that known inward currents can amplify SDs but are not essential for initiating them, supporting an alternative mechanism of SD initiation.

Our findings support the possibility that SDs are mediated by a novel “SD channel” or transmembrane complex, as has been hypothesized (Aitken et al., 1998; Basarsky et al., 1999; Chen et al., 2017; Ricks et al., 2025; Wang et al., 2024). Potential candidates include pannexins (Chen et al., 2017), volume-regulated anion channels (VRACs) (Basarsky et al., 1999; Ricks et al., 2025), or unidentified stretch- or volume-sensitive conductances. The *ex vivo* zebrafish model is well-suited for testing these hypotheses through genetic screening, live imaging, and CRISPR-based mutagenesis.

This study has several limitations. First, we examined SD induction but not propagation across tissue. It is possible that while SDs could still be initiated under the full pharmacological blockade, their spread may have been significantly inhibited, an important question for future studies. Second, the pharmacokinetics of the inhibitory compounds must be considered. Although compounds were applied at effective concentrations to block their respective targets, incomplete tissue penetration or distribution may have allowed residual current. However, consistent results across species and preparations make this unlikely to fully explain the robust SD persistence observed.

Collectively, our findings support a revised conceptual model of SD initiation, centered on the opening of an unknown large-conductance channel rather than classical excitatory ion channel overactivation. The *ex vivo* zebrafish brain preparation introduced here provides a powerful and versatile system for testing these hypotheses in future studies and uncovering the mechanisms underlying SD induction.

## AUTHOR CONTRIBUTIONS

PW: Writing-original draft, Writing-review and editing, Investigation, Validation, Software, Formal Analysis, Data curation, Visualization; AJ: Writing-original draft, Investigation; KA: Writing-review and editing; BC: Methodology, Writing-review and editing; HM: Conceptualization, Methodology, Investigation, Formal Analysis, Visualization; LS: Investigation; Writing-review and editing, Visualization DN: Investigation, Writing-review and editing; JN: Formal Analysis, Writing-review and editing; RA: Investigation; BB: Resources, Project Administration; AS: Conceptualization, Resources, Writing-review and editing; RP: Writing-orginal draft, Conceptualization, Funding acquisition, Resources, Writing-review and editing, Supervision, Project Administration

## ACKNOWLEDGEMENTS

We thank Jordan T. Yorgason and Juha Voipio for their generous donations of lab equipment that made this work possible.

